# Comparing *Caenorhabditis elegans* gentle and harsh touch response behavior using a multiplexed hydraulic microfluidic device

**DOI:** 10.1101/162990

**Authors:** Patrick D. McClanahan, Joyce H. Xu, Christopher Fang-Yen

## Abstract

The roundworm *Caenorhabditis elegans* is an important model system for understanding the genetics and physiology of touch. Classical assays for *C. elegans* touch, which involve manually touching the animal with a probe and observing its response, are limited by their low throughput and qualitative nature. We developed a microfluidic device in which several dozen animals are subject to spatially localized mechanical stimuli with variable amplitude. The device contains 64 sinusoidal channels through which worms crawl, and hydraulic valves that deliver touch stimuli to the worms. We used this assay to characterize the behavioral responses to gentle touch stimuli and the less well studied harsh (nociceptive) touch stimuli. First, we measured the relative response thresholds of gentle and harsh touch. Next, we quantified differences in the receptive fields between wild type worms and a mutant with non-functioning posterior touch receptor neurons. We showed that under gentle touch the receptive field of the anterior touch receptor neurons extends into the posterior half of the body. Finally, we found that the behavioral response to gentle touch does not depend on the locomotion of the animal immediately prior to the stimulus, but does depend on the location of the previous touch. Responses to harsh touch, on the other hand, did not depend on either previous velocity or stimulus location. Differences in gentle and harsh touch response characteristics may reflect the different innervation of the respective mechanosensory cells. Our assay will facilitate studies of mechanosensation, sensory adaptation, and nociception.

## Introduction

The sense of touch allows animals to detect and react to forces resulting from physical contact with the outside world. Much of the pioneering work in identifying the molecules and mechanisms underlying touch response has been done in small genetic model systems such as the roundworm *C. elegans*^1^. This organism’s simple, well-mapped nervous system, optical transparency, short life cycle, and amenability to genetic manipulation make it an attractive model for understanding the molecular and circuit bases of mechanosensation.

Current, widely used behavioral assays for *C. elegans* touch involve either stroking the animal with a fine hair (“gentle touch”) or prodding it with a platinum pick (“harsh touch”). These types of touch sensation have been shown to be mediated by different subsets of sensory neurons^1–4^.

Sensation of gentle touch to the body is mediated by five touch receptor neurons (TRNs). These are ALM right and ALM left (R/L) and AVM in the anterior half of the body, and PLM(R/L) in the posterior half^1^. PVM is sometimes considered a posterior TRN due to morphological and genetic similarities to the other five, but it has not been shown to mediate or contribute to the gentle touch response^5^.

Gentle touch to the anterior of the body usually results in reverse movement, while gentle touch to the posterior of the body usually results in forward movement. Touches to the middle of the body can elicit either response, and are not usually performed in mechanosensory assays^6^. Mutants that fail to respond normally to gentle touch are called “*mec*” for *mec*hanosensory abnormal^6^. Genetic screens with the gentle touch assay have identified many proteins necessary for mechanotransduction, including the degenerin (DEG) / epithelial sodium channel (ENaC) subunit MEC-4^1^. The gentle touch assay has also been used to investigate the nature and mechanisms of sensory adaptation^7^ and sensitization^8^.

Harsh touch to the body using a platinum wire pick elicits similar behavior to gentle touch, but depends on a distinct set of sensory neurons, in addition to at least some of the gentle TRNs^9^. These include BDU, SDQR, FLP, AQR, and ADE in the anterior, and PVD and PDE in the posterior. Harsh touch response is independent of the gene *mec-4*, and has been shown to involve either TRP-4^4^ or the Deg/ENaC subunits MEC-10 and DEGT-1^10^ in different neurons. Harsh touch is thought to be a form of nociception (detection of harmful stimuli) because its response threshold is on the order of the threshold of physical damage^4^. Like mammalian nociceptors^11^, many of the *C. elegans* harsh touch receptors are polymodal sensory neurons, such as PVD(R/L), a pair of highly branched neurons that send processes throughout the body^3,12^.

The gentle and harsh touch manual assays have two important limitations. First, they are low in throughput, being performed manually on one worm at a time. Second, they are largely qualitative in nature, both in terms of the stimulus delivered and the resulting behavior. Tools with very different shapes and mechanical properties are used to test gentle and harsh touch, making it difficult to compare their relative thresholds. These limitations complicate measurement of subtle differences in sensitivity and location-dependence of touch response behavior.

To partially address these limitations, several alternative *C. elegans* touch assays have been reported. Tapping a agar plate containing worms induces touch response behaviors, which can be observed using machine vision^13,14^. Plate tap has been used to study mechanosensory adaptation^7,15,16^. However, this method lacks spatial selectivity, stimulating both the anterior and posterior TRNs, and has not been reported to elicit the harsh touch response^1,17^. Another approach has been to deliver measurable forces to specific locations on a single worm using a piezoresistive cantilever. This method has been used to explore the biomechanical properties of the worm’s body^18^ and their effects on touch sensitivity^19^, and to develop a biophysical model of mechanotransduction in the touch cells^20^. Another approach is to immobilize a single worm with glue^9^ or a microfluidic trap^21,22^ and use a glass probe or pneumatic indenter to apply direct stimulus while monitoring calcium transients or electrophysiological activity^23^. However, no method to date has combined the application of a localized, tunable, mechanical stimulus with behavioral recording of the responses of many worms at the same time.

Here we report a microfluidic-based touch assay that can deliver spatially localized gentle and harsh touch stimuli to up to several dozen *C. elegans* and quantify their behavior before and after stimuli. Our design integrates concepts from several previous microfluidic devices: (1) an array of channels for imaging a large number of *C. elegans* at once^24^, (2) sinusoidal microfluidic channels and ‘artificial dirt’ post arrays that encourage natural crawling behavior^25^, and (3) pressure-actuated monolithic microfluidic valves^26^ that apply localized touch stimuli.

We sought to characterize and compare aspects of the gentle and harsh touch responses on a quantitative level. First, we measured the relative thresholds for gentle (*mec-4* dependent) and harsh (*mec-4* independent) touch. Next, we investigated the extent of gentle touch receptive field overlap by comparing the receptive fields of wild-type and mutant animals. We then examined the influence of prior behavior and prior touch history on behavior after an ambiguous stimulus (touch to the mid-body), for both gentle and harsh touch.

## Experimental

### Device Concept and Design

Our microfluidic device (Fig. 1) consists of: (1) a layer containing loading channels with six bifurcations leading to an array of 64 sinusoidal channels into which worms are allowed to crawl, and (2) a layer containing an array of 15 channels that can be pressurized to deliver touch stimuli to worms in the first layer. Each intersection between the worm channels and touch channels, which are mutually perpendicular, forms a monolithic microfluidic valve^26^ capable of partially closing the worm channel and delivering a touch stimulus if a worm is present (Fig. 1a). The touch channels are filled with water to minimize compressibility and reduce optical scattering arising from the refractive index difference with the PDMS device.

**Fig. 1.**
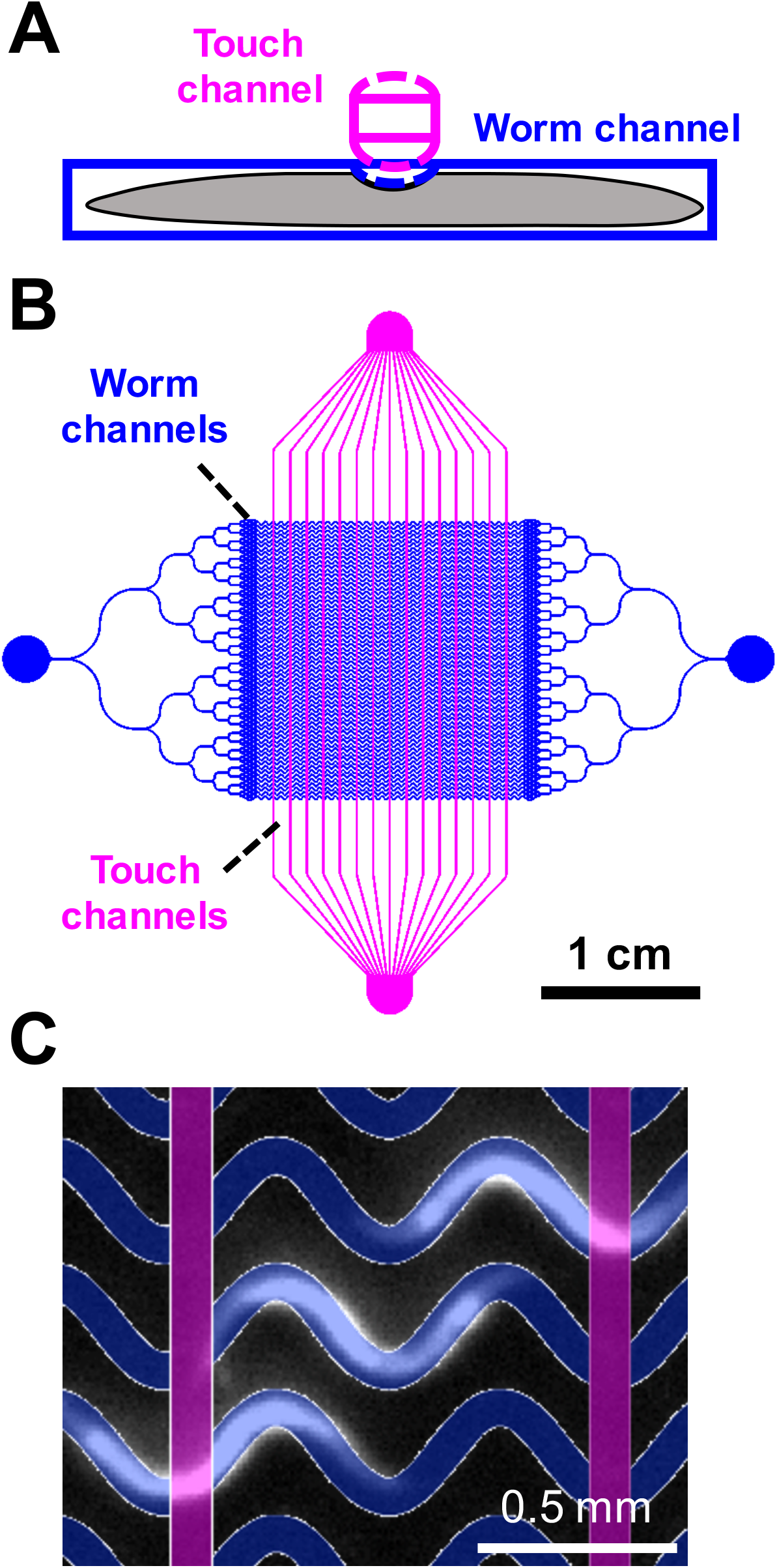
Microfluidic device for assaying touch response behavior, (a) A touch channel (magenta) inflates upon pressurization, partially closing the worm channel (blue), (b) Schematic of the device containing 64 sinusoidal worm channels and 16 control channels, (c) Dark field image of *C. elegans* crawling in the worm channels with photomask design overlaid.

Worms are loaded through an entry port and flow through a set of bifurcating channels that distribute them approximately uniformly^24^ before entering the worm channels. All layers of the device are transparent, allowing for behavioral imaging under dark field or bright field illumination.

The design of our device reflects optimizations over several constraints and trade-offs. The number of channels and the field of view were chosen to accommodate as many animals as possible while allowing sufficient spatial resolution to clearly identify worms including their anterior-posterior orientation. The touch channels were spaced 1.067 mm apart, approximately the length of an adult worm, such that each animal experiences only one stimulus at a time (Fig. 1c). The sinusoidal shape of the worm channels allows the animals to exhibit a natural crawling behavior similar to that on a moist surface^25^, and clearance in the worm channels allows worms to execute turns and pass one another. The thicknesses of the layers was optimized over several iterations to allow repeated gentle or harsh touch stimuli while maintaining the integrity of the device. Thicker separations between the worm and control channels resulted in smaller deflections and the inability to trigger *mec-4*-independent touch response; thinner separations led to device failures after repeated pressure cycles.

### Mold fabrication

We designed photomasks (Supplemental Information S1 and S2) in DraftSight (Dassault Systèms, Vélizy-Villacoublay, France) and had them printed on polyester film by Fine Line Imaging (Colorado Springs, CO). We fabricated worm and control layer molds using standard soft lithography techniques. Briefly, SU-8 2025 (MicroChem, Westborough, MA) was spin-coated onto a 5-inch diameter Si wafer for 10 s at 500 RPM followed by 30 s at 1000 RPM (worm layer) or 500 RPM (control layer). After soft baking at 80 °C for 10 min (worm layer) or 60 °C for 2 h (control layer), wafers were placed under a 360 nm long pass filter and treated with a 2.6 J/cm^2^ (worm layer) or 3.2 J/cm^2^ (control layer) ultraviolet exposure in an Intelli-Ray 400 UV curing oven (Uvitron, West Springfield, MA). We developed photoresists by immersion in propylene glycol monomethyl ether acetate (Sigma-Aldrich). Molds were silanized with trichloro(1H,1H,2H,2H-perfluorooctyl)silane (Sigma-Aldrich) for 20 min to facilitate demolding.

### Device fabrication

The elastic modulus of PDMS can be adjusted by varying the ratio of base to curing agent^27,28^. To create the worm layer, we mixed PDMS (Dow Corning Sylgard 184) at a 20:1 base:curing agent ratio, degassed under vacuum for 30 min, spin-coated it onto the mold for 90 s at 630 RPM, and baked it on a level hotplate at 50 °C overnight (~12 hr). To create the control later, we mixed PDMS at a 5:1 or 10:1 base:curing agent ratio, poured it onto the mold in a petri dish to a depth of 10 mm, vacuum degassed it for 30 min, and cured it in a 50 °C oven overnight.

To bond device layers, we plasma treated the surfaces to be bonded for 9 s in a plasma cleaner consisting of a Plasma Preen II 973 controller (Plasmatic Systems, Inc.) connected to a modified microwave oven (Amana RCS10TS) and then pressed the surfaces together for several minutes. We first demolded the control layer and bonded it to the worm layer. Next we demolded both layers from the worm layer mold and plasma bonded the worm layer side to a 75 mm × 25 mm × 1 mm glass slide. Each device was calibrated before use (see Results).

### Control system

Control pressures were provided by a nitrogen gas cylinder through a two-stage pressure regulator (Harris Products) and measured by an analog pressure gauge. We used a 3-way solenoid valve (Asco 3UL87) to apply or release pressure to the touch channels. The solenoid valve was controlled with the analog output of a National Instruments USB-6001 DAQ device coupled to a solid-state relay.

### Imaging system

We recorded behavior at 10 frames per second with a 5 megapixel CMOS camera (DMK 33GP031, The Imaging Source, Charlotte, NC) and a C-mount lens (Schneider Kreuznach Xenoplan 1.4/23-0512, 23 mm effective focal length) using IC Capture software (The Imaging Source) on a Windows PC. The field of view was approximately 15 mm × 11 mm. To ensure sufficient resolution for tracking, we did not image all 64 worm channels at once. Red LED strips (Oznium, Inc.) surrounding the device provided dark field illumination.

### Experimental procedures

To prepare the device, we first filled the control channels and connecting tubes with water. Then we filled the worm channels with NGMB (50 mM NaCl, 1 mM CaCl_2_, 1 mM MgSO_4_, 20 mM KH_2_PO_4_, 5 mM K_2_HPO_4_) containing 0.1% bovine serum albumin (Sigma A9647) to minimize adhesion of worms to the channels and tubing. We used NGMB to wash *C. elegans* from their growth plates and placed them in a syringe connected to the worm channel inlet port. To load worms into the device, we used syringes on the inlet and outlet tubes to manually apply pressure or vacuum. Loading takes approximately 5 minutes. All experiments were performed at room temperature (18-22 °C).

For each set of worms, we first recorded for at least 30 s to establish a baseline level of behavior. Next, we applied one sham stimulus with zero pressure followed by one of two stimulus regimes. (1) To determine the response threshold of a population of animals, we delivered a ramp of twelve stimuli of increasing magnitude with a 30 s inter-stimulus interval (ISI). (2) To determine the sensory adaptation and / or behavioral receptive field of a population of animals, we delivered a series of 20 equal magnitude stimuli with a 30 s ISI. Data collection time was 7 min and 11 min, respectively. For all worm experiments, each stimulus consisted of a train of 5 pulses of 20 ms duration with 20 ms separation between pulses. Supplemental Video S3 shows a subset of worms on the device undergoing a touch stimulus.

After each experiment, the device was cleared of worms by flowing a bleach solution (1:1:3 parts by volume mixture of 5 M NaOH, 5% NaClO, and water) through the worm channels for approximately 5 min., followed by a 5 min water rinse and refilling with NGMB. Thus a typical 20-stimulus experiment lasted about 25 minutes and involved 50 worms. The overall experimental throughput was approximately 2400 individual touch assays on 120 distinct worms per hour.

### *C. elegans* strains

Strains used in this study were Bristol N2 (WT), TU253 *mec-4(u253)*, and TU4032 *egl-5(u202); uIs115 [Pmec-17::RFP]*. Animals were cultured on OP50 *E. coli* food bacteria on standard NGM agar plates^13^ or high-peptone NGM plates (same as NGM plates except with 10 g/L peptone) at 15-20 °C. To synchronize growth, we used a sodium hypochlorite bleach procedure^29^ to obtain eggs, which were hatched in NGMB overnight. About 200 worms were then transferred onto OP50 seeded NGM agar plates and grown to adulthood. All experiments were performed using day 1 adult hermaphrodites.

### Image processing

All image processing and data analysis was performed using custom software written in MATLAB (MathWorks, Natick, MA). Briefly, each frame was background subtracted and thresholded to obtain a binary image of the worms on the dark background. We determined the head-tail orientation of each worm by visual inspection. Velocities were calculated by tracking the centroid of each animal over time.

We excluded images acquired during each stimulus because valve actuation caused a small distortion in the device and a fluctuation in animal position. The worm channels provide enough clearance for worms to pass each other or execute a 180 degree turn (see Supplemental Video S4). We excluded from analysis worms that were touching or overlapping. We also excluded worms that were turning because they could receive touch stimuli to two locations simultaneously.

## Results and Discussion

### Stimulus calibration and measurement

One limitation of traditional touch assays in *C. elegans* is the difficulty of controlling the strength of stimulus delivered to the animal by hand. In our device, stimulus amplitude can be continuously varied by changing the pressure delivered to the touch channels, causing the worm channel ceiling to deflect downward by variable amounts. The microvalve indenter is slightly rounded when pressure is applied. As for any rounded indenter (*e.g*. eyebrow hairs, wires, glass probes, and microspheres^6^) the contact area between indenter and worm increases with indentation depth.

Previous work^19,20^ showed that the amount of deformation, not pressure, is the key determining quantity for the mechanoreceptor response. We therefore used deformation amplitude as the measure of stimulus amplitude. To calibrate the relationship between pressure and deformation of each device, we measured the worm channel height inside the microvalves at different pressures by monitoring the transmission of light through a blue dye solution (Fig. 2). We filled the worm channels with 15 mM Brilliant Blue FCF dye in water and recorded video sequences of valve closure at different pressures under bright field illumination provided by a red LED (Fig. 2a). The Beer-Lambert Law describes the relationship between the proportion of light transmitted through the channel (I/I_0_) to the worm channel height (L): I/I_0_ = exp(-L/λ) + B, where λ is the absorption length and B is the baseline intensity when transmitted light is blocked. We used the known height of the worm channel when fully open (75 μm) and the intensity of a region of interest (ROI) in which all transmitted light was blocked with an opaque material to calculate λ and B, respectively. We used this relationship to convert the light intensity recorded in an ROI to the deflection of the worm channel ceiling in the touch valves (Fig. 2b). For calibration experiments, we used trains of 10-20 pulses to compensate for the video’s sparse sampling of the device response. We repeated this procedure at least three times to develop a calibration curve for each device (Fig. 2c). By testing several valves at the center and edges of a single chip, we verified that calibrations were uniform throughout the device. Thereafter we calibrated each device using a single valve at the center of the chip.

**Fig. 2.**
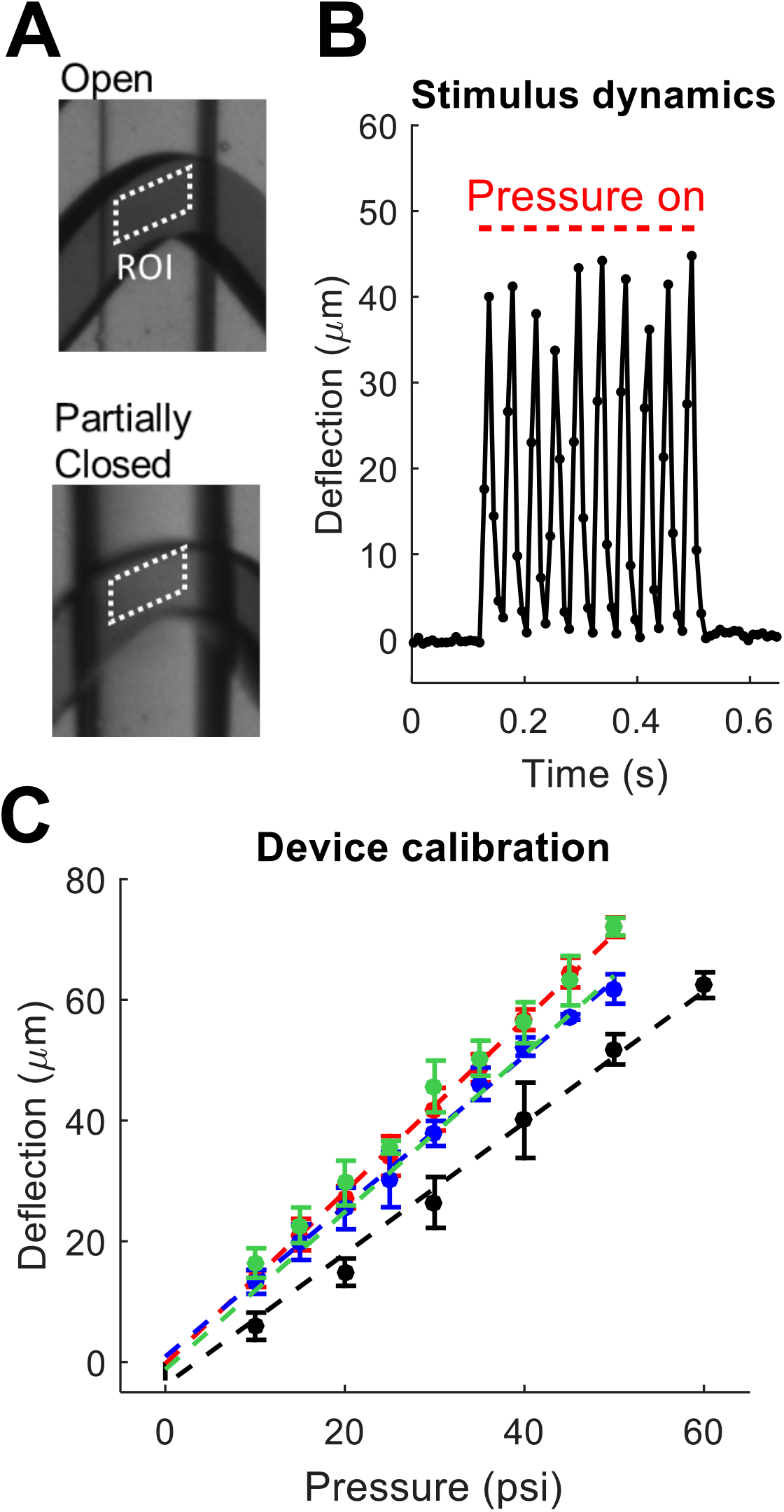
Stimulus measurement and calibration. (a) Optical transmission of dye-filled touch valves is used to monitor worm channel height in the valve. (b) Worm channel ceiling deflection when control solenoid is driven by a 25 Hz square wave with an amplitude of 40 psi and a 50% duty cycle. Red lines denote pressure on. (c) Maximum deflection as a function of pressure. Points (mean ± SD) show the average of 3-5 trials. Colors show 4 different devices.

The calibration procedure could not be performed with a worm present in the valve because the worm’s body excludes the dye and blocks some of the transmitted light. The values given for deflection are therefore for a worm channel not containing a worm. To determine to what extent the presence of an animal changes the touch valve deflection, we measured the channel deflection using a confocal microscope (Leica SP5). We loaded day 1 adults into the device and immobilized them in a solution of 10 mM NaN_3_ with 0.3 μM sodium fluorescein in NGMB. We imaged the 3-dimensional shape of the fluorescein solution in two adjacent touch valves, one containing a worm and the other containing only the fluorescein solution. Due to the long acquisition time required for confocal microscopy, we applied a static instead of pulsatile pressure. We used 15 psi because maintaining higher pressures for several minutes compromised the device integrity. We found no difference between the deflection of the touch valves with a worm (deflection 19.1 ± 2.0 μm (mean ± SD) at 15 psi) and without a worm (deflection 19.4 ± 1.7 μm at 15 psi). Both values agreed with results from optical transmission studies described above (deflection 18.8 ± 0.3 μm at 15 psi).

We conclude that the presence of a worm does not significantly affect the deflection of the microfluidic valve at this pressure. This may be because the worm’s elastic modulus is much smaller than that of PDMS. Studies that consider the worm as a whole have estimated its modulus to be the range of 110-140 kPa^30,31^, compared to ~1 MPa for PDMS with a 20:1 base:curing agent ratio^32^. However, the stiffness of *C. elegans* and other biological tissues is known to increase sharply with strain^31^, so it is possible that valve deflection with and without a worm are not equal at higher pressures.

### Comparison with classical touch assays

We sought to determine to what extent the touch response behavior in our device is similar to that on an agar plate. We performed the traditional (eyebrow hair) anterior gentle touch assay on 10 worms crawling on an unseeded agar plate while acquiring video recordings on a stereo microscope. We measured the wavelength and bending frequency of the animals in 3 second windows before and after the touch. We found no change in wavelength (0.49 ± 0.4 mm before and 0.51 ± 0.3 mm after, p = 0.22, 2 tailed paired t-test) but a significant increase in frequency 0.59 ± 0.31 Hz to 1.39 ± 0.38 Hz (p = 9.2 * 10^-6^, 2-tailed paired t-test). In the microfluidic chip, we also saw a significant increase in frequency in the three seconds after the stimulus, from 1.00 ± 0.58 Hz before stimulus to 2.56 ± 0.88 Hz after stimulus (p = 6.6 * 10^-5^). The worm’s wavelength in the microfluidic device was constrained to be 0.5 mm, very close to that observed on agar. While the worms move faster overall in the microfluidic channels, both worms on plates and in our device respond to touch by increasing their bending frequency by similar amounts (2.36 times on agar, 2.56 times in the microfluidic device). These results show that with regard to touch response behavioral characteristics, the microfluidic device environment is reasonably similar to that of an agar plate.

### Comparison with other quantitative *C. elegans* touch assays

Two classes of existing assays allow for tunable touch stimuli, as well as quantitative response data. The first class is based around the plate tap reflex and comprises assays that use an impactor or actuator to induce vibration of the agar substrate, triggering a response mediated by the gentle touch receptors^14^. As in our assay, stimulus strength can be measured, usually with a MEMS accelerometer^7,33,34^ or laser Doppler vibrometer^35^, and responses of freely moving animals can be recorded. However, the stimulus is not localized to any one part of the animal, preventing the study of touches to a subset of touch receptors.

The second class involves directly touching a single, often immobilized animal with instruments such as glass micropipettes, piezoresistive cantilevers, and microfluidic actuators. A previously reported microfluidic actuator^21^ with an in-plane deflection geometry has a smaller standard deviation of deflection (1 μm) than our device (2.6 μm). However, our device has a two-layer geometry that permits assaying many animals simultaneously. Thus our assay combines the multi-worm throughput and quantitative behavior measurement of the substrate vibration assays with the localized, tunable stimuli of the direct touch assays.

### Quantification of gentle and harsh touch response thresholds

A quantitative understanding of touch response behavior is necessary for understanding which cells and genes are required to detect different physical properties and govern different aspects of the response. In traditional touch assays, gentle and harsh touch are assayed using different tools, and animals are normally scored in a binary fashion as responding or not responding. However, touch responses are known to vary both qualitatively (*e.g*. direction of movement) and quantitatively (distance travelled during response)^4,14^.

Worms lacking the DEG/ENaC channel subunit MEC-4 are insensitive to gentle touch but remain sensitive to harsh touch^4^. While some estimates of the forces and/or deformations required for gentle and harsh touch have been reported^4,19,23,36^, these measurements have been performed in different ways, for example using unequal probe sizes, making them difficult to compare directly. We used our touch microfluidic device to measure the differences in touch sensitivity between N2 and *mec-4* worms.

To determine the threshold (defined here as the stimulus amplitude at which the response probability is 50%), we delivered stimuli of monotonically increasing strength spaced 30 s apart. Since response to local touch occurs quickly, we quantified the worms’ behavioral responses by measuring the difference in centroid velocity of the animals between one second before and one second after the stimulus (Fig. 3). To determine a threshold for automatic scoring, we scored a subset of our data (149 worm touches) by visual inspection of video recordings and then chose the threshold (0.18 mm/s) that maximized the difference between the true positive rate (80% for this threshold) and the false positive rate (14.7% for this threshold). Animals whose velocity changed more than the threshold after the stimulus were scored as responding. We excluded animals that received a posterior touch if they were already moving forward, as well as animals that received an anterior touch if they were already reversing. We adjusted for possible false positives by subtracting the response rate of the strain to a mock stimulus (12.3% for WT, 0% for *mec-4*). We considered the lowest deflection causing at least 50% of animals tested to respond to be the response threshold, interpolating as necessary. For WT animals, the threshold was 9.3 ± 2.7 μm, and for *mec-4* animals, the threshold was 46.1 ± 2.8 μm. For WT animals, response probability decreased for the strongest stimuli, suggesting that a MEC-4-dependent sensory adaptation or response fatigue occurs with increasing mechanical stimulus amplitude.

**Fig. 3.**
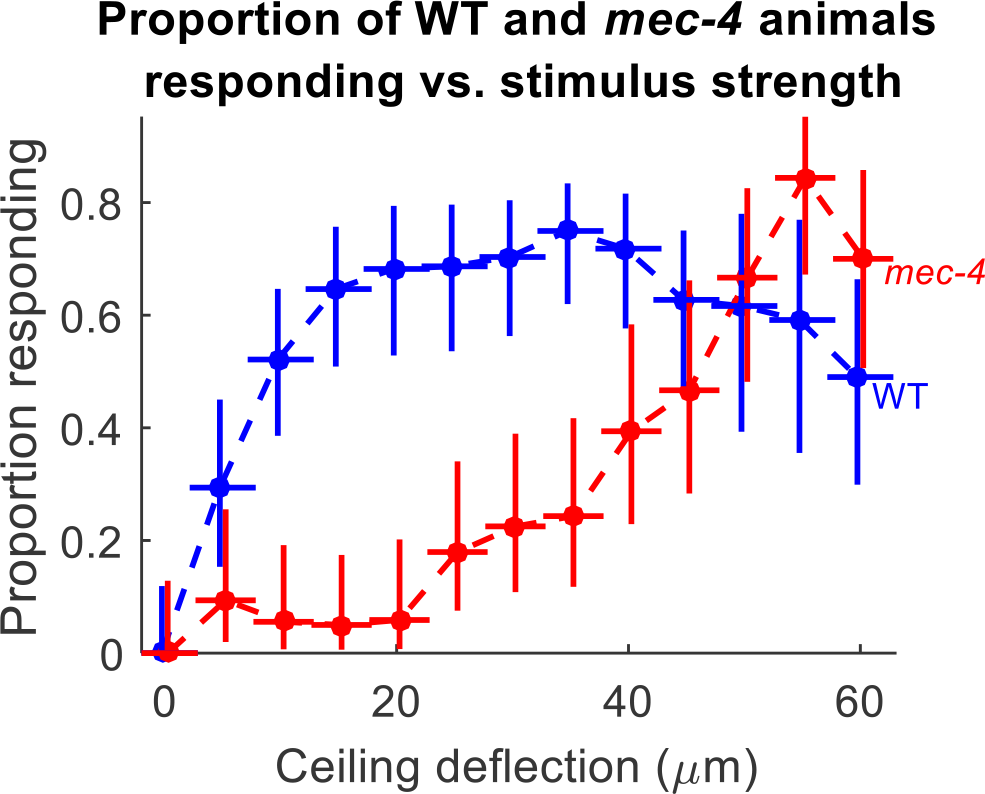
Proportion responding of WT and *mec-4* as a function of stimulus amplitude. Vertical error bars are 95% confidence intervals based on a binomial fit, and horizontal error bars are standard deviations of the calibration measurements for the device. Each point represents data from n=21-57 animals.

These results demonstrate the ability of this device to administer a continuum of mechanical stimuli, and show that in terms of the vertical deflection of the microfluidic membrane in this assay, the threshold for harsh touch is about 5 times greater than the threshold for gentle touch.

### Estimates of the force applied to the worm

Recent studies suggest that touch receptor neurons are sensitive to deformation rather than force *per se*^19,20^. Nevertheless, touch stimuli are often quantified in terms of force applied^6,23^. While we have no direct measure of the force applied to the worm in our device, we can approximate this force based on the deflection and previous estimates for the mechanical properties of the worm. Our confocal measurements suggest that touch valve deflection is similar with and without a worm up to at least 15 psi, the pressure tested. A deflection of 9.3 μm causes a response in WT animals 50% of the time, and the Young’s modulus of the worm, based on an average of literature values, is approximately E=125 kPa (average of two estimates^30,31^). Since the worm’s modulus is much smaller than that of PDMS, the force on the worm is dominated the worm’s elasticity. A simple estimate based on the elasticity of the worm yields yields a force of 116 μN for a 9.3 μm deformation. Application of deformation theory^37^ for a cylindrical body in contact with a surface of limited extent yields an estimate of 94 μN (Supplemental Information S5).

Using 10 μm diameter glass bead as an indenter, Petzold *et al*. reported a 50% response probability at a force of about 0.75 μN and deflection of 1.5 μm, both much smaller than we find here. This difference is likely due to the very different sizes of the indenters, and differences in indentation number, rate, and shape. If we consider the active area of the spherical indenter to be the area of a disk with the same diameter, and the active area of a touch valve to be a rectangle (Supplemental Information S5) the downward force per area exerted by the touch valve and glass bead are similar: 19.0 kPa and 18.7 kPa, respectively.

### Behavioral receptive fields of the touch receptor neurons

*C. elegans* normally responds to touch by moving in a direction opposite to the stimulus: anterior touch results in a reversal and posterior touch results in forward movement. Touches near the middle of the body can result in either forward or reverse movement, and Ca^2+^ imaging of ALM shows sporadic activation by near (close to mid-body) but not far posterior touch^9^. This suggests an overlap in receptive fields between the anterior and posterior TRNs, even though the TRN processes themselves do not overlap. However, the receptive fields, or regions over which the anterior and posterior are sensitive to touch, have remained unclear.

To spatially map the gentle touch behavioral response, we used the ability of our assay to rapidly assess behavioral responses as a function of body position. Using our microfluidic device, we measured the responses of WT animals and *egl-5(u202)* mutants, in which the posterior TRNs are not functional^38^. For each group of worms, we administered 20 stimuli with a deflection amplitude of 30 μm separated by a 30 second ISI and measured the speed for one second after the stimulus. The average speed after a stimulus was 0.35 mm/s for the first five stimuli and 0.30 mm/s for the final five stimuli in WT animals, indicating that sensory habituation is minimal in this protocol (Supplemental Fig. S6). The average speed was 0.21 mm/s after a mock stimulus.

*egl-5* mutants were slower than WT, *egl-5* worms in the absence of stimulus had an average speed of 0.10 mm/s versus 0.23 mm/s for WT, and also had a significantly slower response speed (p < 0.05, two-sided t-test), but their responses also did not decline significantly over the course of the experiment.

To examine responses to touches at different regions of the body, we collected the responses into five bins according to body coordinate touched (Fig. 4). We excluded data from touches to the anterior and posterior 15% of the body because of reduced responsiveness to touches in these region, possibly due to decreasing body diameter at the two ends of the animal, which could reduce the stimulus experienced. The rest of the body also varies in width (Fig. S7), so we do not make direct comparisons between different location bins.

**Fig. 4.**
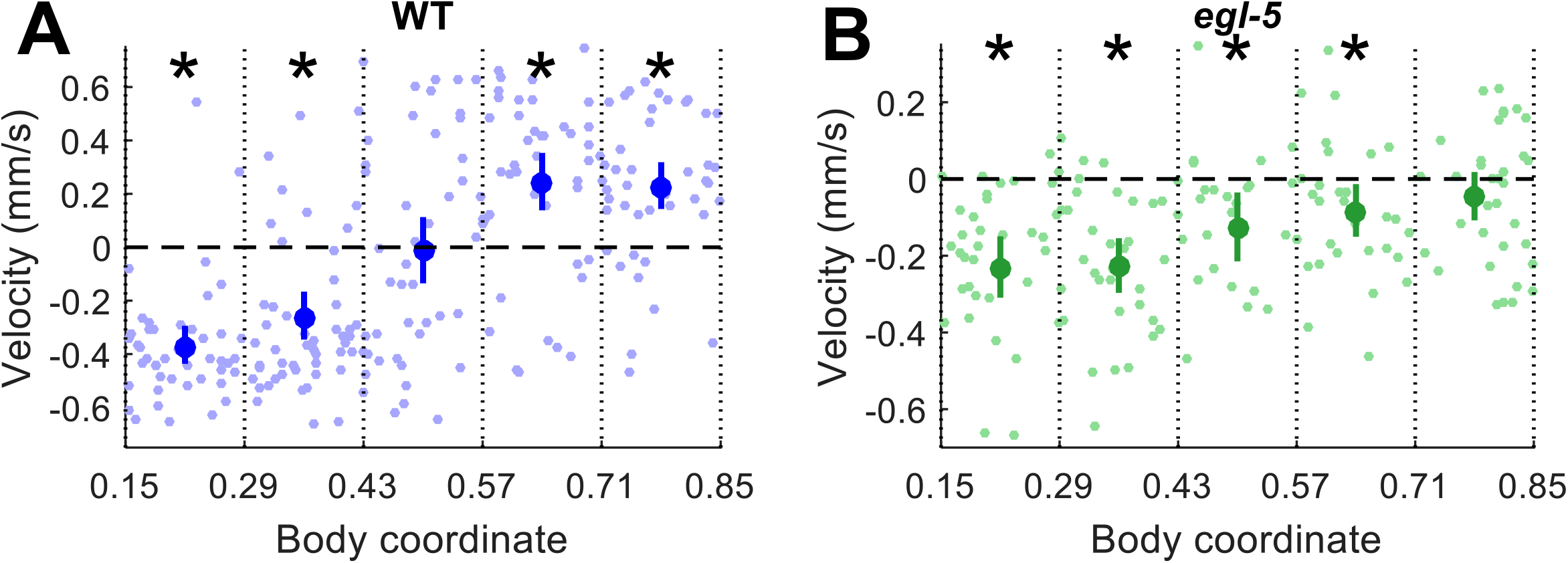
Gentle touch response fields. For both plots, responses are grouped into five bins by body coordinate (0 = head, 1 = tail) of the center of stimulus. Average ± 95% CI of the mean is plotted for each bin. Asterisks denote significant difference from zero for the responses in the bin (Z test, p < 0.05 after Bonferroni correction). Pale dots show responses of individual animals. (a) Velocity of WT animals after gentle touch stimuli. (b) Velocity of *egl-5(u202)* animals after gentle touch stimuli.

WT animals responded to anterior touch by movement in the reverse direction (negative mean velocity) and to posterior touch by movement in the forward direction (positive mean velocity) (Fig. 4a). Touch near the center of the worm induced either forward or reverse movement with roughly equal probability (mean velocity close to zero). Variation in response direction decreased for touches close to either end of the animal. These results are consistent with *C. elegans* behavior on agar plates and show that worms’ normal mechanosensory behaviors are retained in our microfluidic device.

Like WT animals, *egl-5* mutants responded to anterior touch by movement in the reverse direction. However, *egl-5* worms also responded to mid-body and mid-posterior touch by reversal, reflecting the absence of posterior TRN function. Responses after mid-body and mid-posterior touch were weaker than that of anterior touch, and, in the most posterior bin, responses were no longer statistically significantly different from zero (p = 0.49, one-sided z-test). *mec-4* worms lacking all TRN function did not respond to touch at this location with reversals (Supplemental Fig. S8), showing that *egl-5* reversals in response to posterior touch are mediated by the anterior TRNs.

Our results show that the anterior gentle touch receptor neurons, in addition to being sensitive to touch in the anterior half of the animal, are also sensitive to touch to the posterior of the body. That is, the receptive fields of one or more of the anterior TRNs extend into the animal’s posterior half. This result suggests that TRNs respond to both local and global touch deformation^19^ through mechanical coupling.

### Influence of previous locomotory direction on touch response

Because the anterior and posterior TRNs influence the motor behavior in an opposing manner, the behavioral response after a touch at or near the middle of the body represents a decision between two conflicting inputs. We examined factors influencing the behavioral response to mid-body touch.

One such potential factor is the worm’s direction of movement prior to the stimulus. It is not clear to what extent the worm’s locomotory behavior before the stimulus influences the behavior after the stimulus. One possibility is that a mechanosensory stimulus induces a certain change in velocity independent of the original velocity, such that the final (after stimulus) velocity is linearly related to the initial velocity with unity slope. A second possibility is that the final velocity is unrelated to the initial velocity. A third, intermediate, possibility is that the final velocity depends on the initial velocity, but with slope less than 1.

To determine the relationship between the velocities before and after each stimulus, we delivered gentle touch stimuli with 30 μm deflection amplitude to WT animals (Fig. 5a). We found that the velocity prior to stimulus had very little influence on the velocity after stimulus. The initial velocity could explain only 4.1% of the variance in the final velocity, while the position of stimulus could explain 32% of the variance in the final velocity. Furthermore, when we grouped responses according to whether the touch stimulus occurred in the anterior, middle, or posterior third of the body (roughly the regions covered by the anterior TRNs, all TRNs, and posterior TRNs) and then compared responses when the worm was initially moving forward to responses when the worm was initially moving backward, we did not find a significant difference in velocity after stimulus (p = 0.32, 0.72, and 0.19 for anterior, middle, and posterior touches, respectively, two-tailed t-test). We also performed the same experiment for harsh touch, using a 50 μm stimulus on *mec-4* animals, and found similar results (Fig. 5b).

**Fig. 5.**
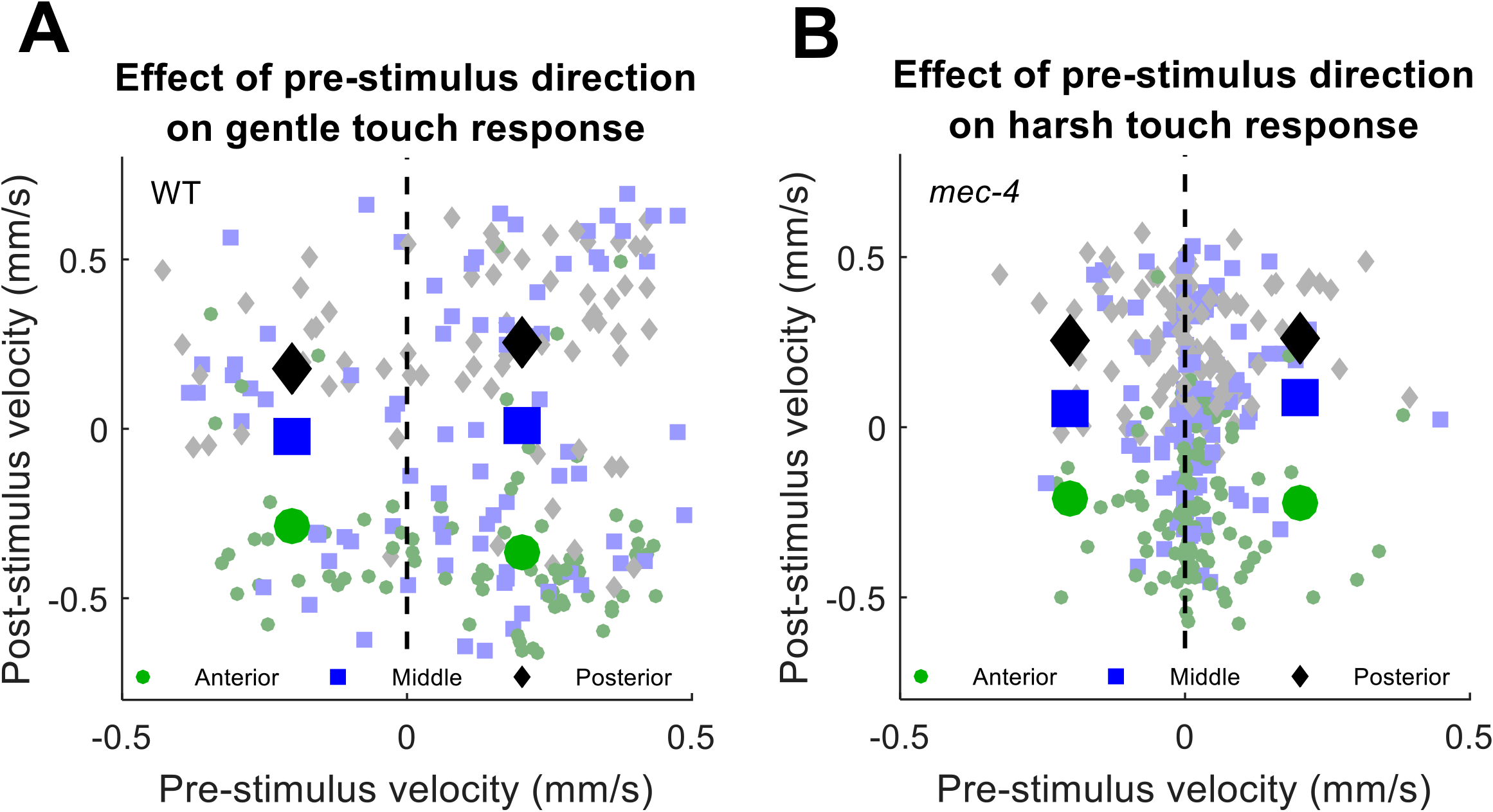
Relationship between pre-stimulus velocity and post stimulus velocity for gentle touch in WT worms (a) and harsh touch in *mec-4* worms (b). Responses are classified by touch location (green circles = anterior, blue squares = middle, black diamonds = posterior). Large shapes represent averages of animals moving backward or forward prior to the stimulus. Small tinted shapes represent individual animals in the same categories. In no case does velocity prior to the stimulus have a significant effect on velocity after the stimulus (p > 0.05, two-tailed t-test with Bonferroni correction)

Our finding that the final velocity is unrelated to the initial velocity suggests that the mechanosensory stimulus applied during gentle or harsh touch to the body resets the forward/reverse state of the locomotory network such that the prior state of the locomotory interneurons does not influence the response direction after gentle touch. This is true even for touches to the middle of the body, where one might expect that balanced inputs from the anterior and posterior TRNs would not upset the bistable^39^ locomotory interneuron network.

This finding is consistent with a rate equation model, such as one described by Roberts et al ^39^ in which reciprocal connections between forward and reverse subcircuits mediate stochastic fluctuations between the respective behaviors, and the amount of forward or reverse circuit activity is boosted by a mechanosensory input.

### Influence of previous touch location on touch response

Next, we asked to what extent previous touches influence the touch response behavior. Such influences may occur due to habituation of the touch response in a position-dependent manner. We conducted gentle and harsh touch experiments as above, with 20 gentle (30 μm deflection) or harsh (50 μm deflection) touch stimuli and a 30 s ISI. We restricted our analysis to animals tracked continuously through two consecutive stimuli. We then grouped the responses to the second touch by the location of the first touch for comparison (Fig. 6).

**Fig. 6.**
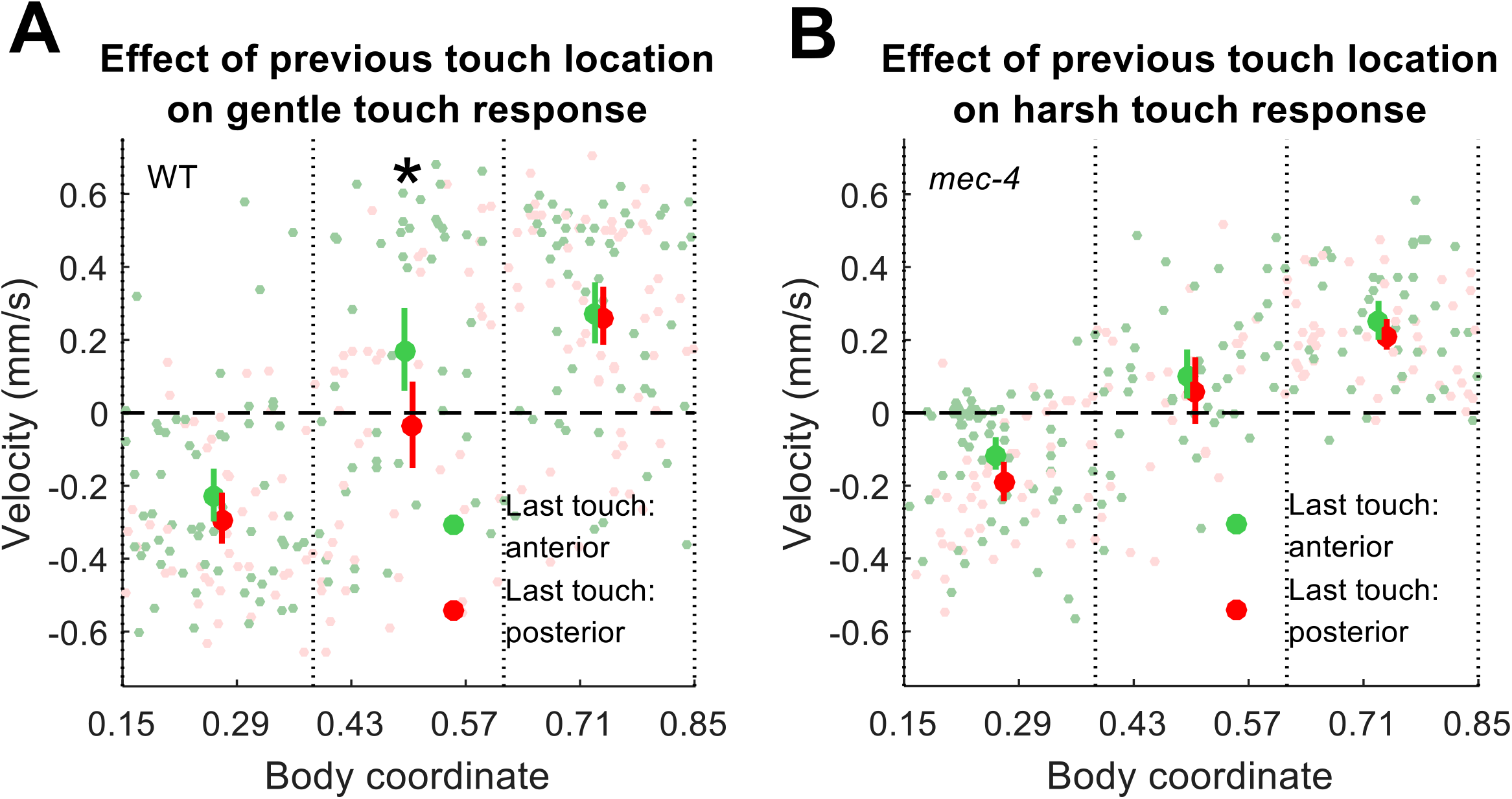
Relationship between preceding stimulus location and post-stimulus velocity for gentle touch in WT worms (a) and harsh touch in *mec-4* worms (b). Responses are grouped into three bins by body coordinate of the current touch (0 = head, 1 = tail) and colored by the location of the preceding touch (green = previous touch to the anterior half, red = previous touch to the posterior half). 95% CI of the mean is shown for each bin. Pale dots show responses of individual animals. The asterisk denotes a significant difference between the two groups in the location bin (two-tailed t-test, p = 0.0315 after Bonferroni correction).

For gentle touches to the middle third, but not the ends, of the body, we found that a previous touch to the anterior half of the body significantly shifted responses in the forward direction compared to a previous touch to the posterior half of the body (Fig. 6a). The simplest explanation of this result is that an anterior touch causes adaptation in the anterior TRNs, changing the balance of sensory input from touches to areas where the receptive fields overlap.

In contrast, when this experiment was done in *mec-4* mutant animals with stimulus amplitudes corresponding to harsh touch, we did not observe any effect of previous touch location on touch response (Fig 6b). This is not due to a lack of a decrement in responsiveness, as overall responsiveness to harsh touch decreases more rapidly with subsequent stimuli than for gentle touch (Fig. S6b). Our results suggest that harsh touch habituation, unlike gentle touch habituation, does not occur in an anterior/posterior-dependent manner. Our result is consistent with an organism wide rather than local regulation of harsh touch.

## Conclusions

We have presented a microfluidic-based method for delivering continuously variable, localized touch stimuli to many freely behaving *C. elegans*. First, we measured the relative response thresholds of gentle (*mec-4* dependent) and harsh (*mec-4* independent) touch, establishing our ability to test both modes of touch with a single assay. This ability, combined with the amenability of microfluidics to the administration of chemical, pharmacological, and optical stimuli, opens the door to studies of the relationship between touch and nociception in *C. elegans*.

Using the ability of our device to provide localized stimuli to many animals simultaneously, we mapped the position-dependent behavioral responses of WT animals and mutants lacking function in some or all of the TRNs. By comparing these responses, we showed that the mutants respond to posterior touch by reversing, showing that the behavioral receptive field of the anterior touch receptor neurons extends into the posterior half of the body. This sensitivity to nonlocal deformations may occur via biomechanical coupling of induced strain through the worm’s body^20^. Together, these experiments demonstrate the utility of our methods for studying how touch response thresholds vary, how they adapt to repeated stimuli, and the extension of a neuron’s receptive field by body mechanics.

Finally, we used our assay to ask what determines the behavioral decision of a worm subject to local touch stimulus. We found that there is little to no influence of the pre-stimulus velocity on the response velocity, for either gentle or harsh touch stimuli. This result supports a model in which the mechanosensory stimulus resets the locomotory network state.

We found a significant effect of the location of the previous gentle touch on responses to gentle touches to the middle of the body, suggesting that touch sensitivity is locally regulated in response to a localized touch. This result is consistent with previous studies showing that the anterior and posterior gentle touch circuits can be modulated independently^7,17,40^. However, no such location-dependent sensitivity to previous touch was observed for harsh touch stimuli. This difference may reflect the whole-body innervation of some of the harsh touch receptor neurons, as compared to the anterior and posterior specific innervation of the TRNs mediating gentle touch.

Our findings open new possibilities in generating and testing quantitative models of the touch response and the coupling between the mechanosensory and the locomotory networks.

## Acknowledgements

We thank David Issadore for technical support and equipment and Martin Chalfie for strains. Some strains used in this study were provided by the CGC, which is funded by the NIH Office of Research Infrastructure Programs (P40 OD010440). P. D. M. was supported by the National Institutes of Health. J. H. X. was supported by the Center for Undergraduate Research and Fellowships. C. F.-Y. was supported by the National Institutes of Health, Ellison Medical Foundation, and Sloan Research Foundation.

## Supplementary Information

**S1:**
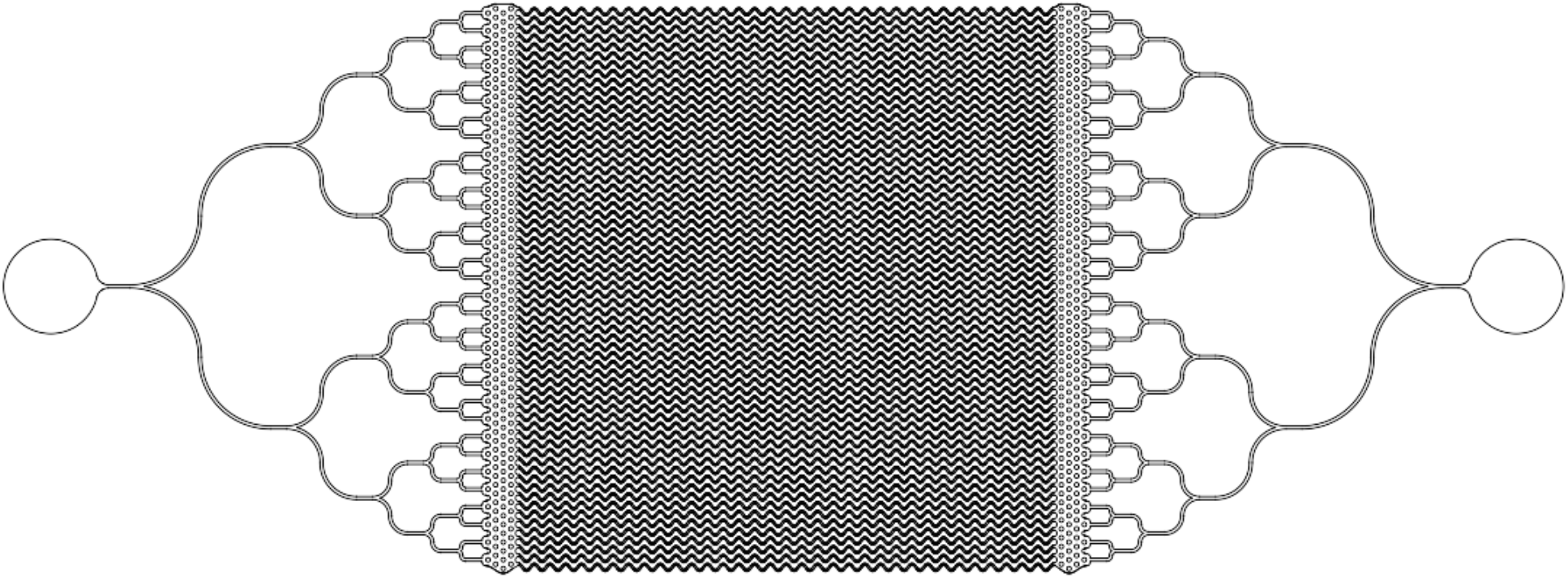
Photomask for the worm layer (DWG file)

**S2:**
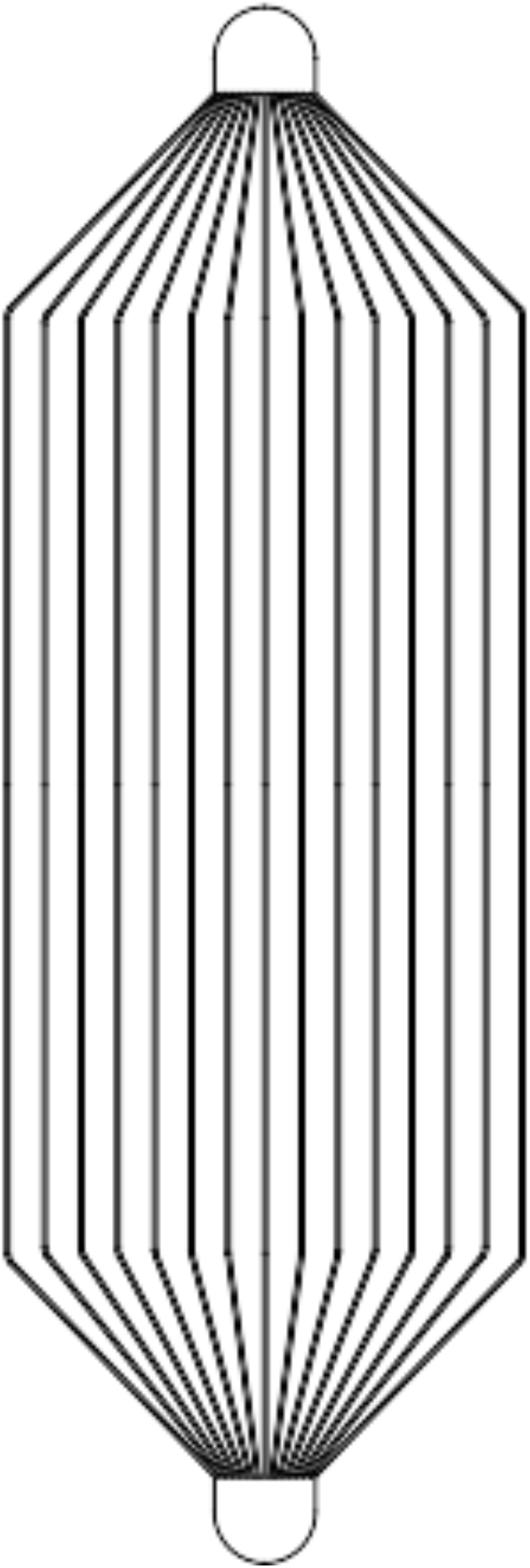
Photomask for the control layer (DWG file)

**Video S3:**
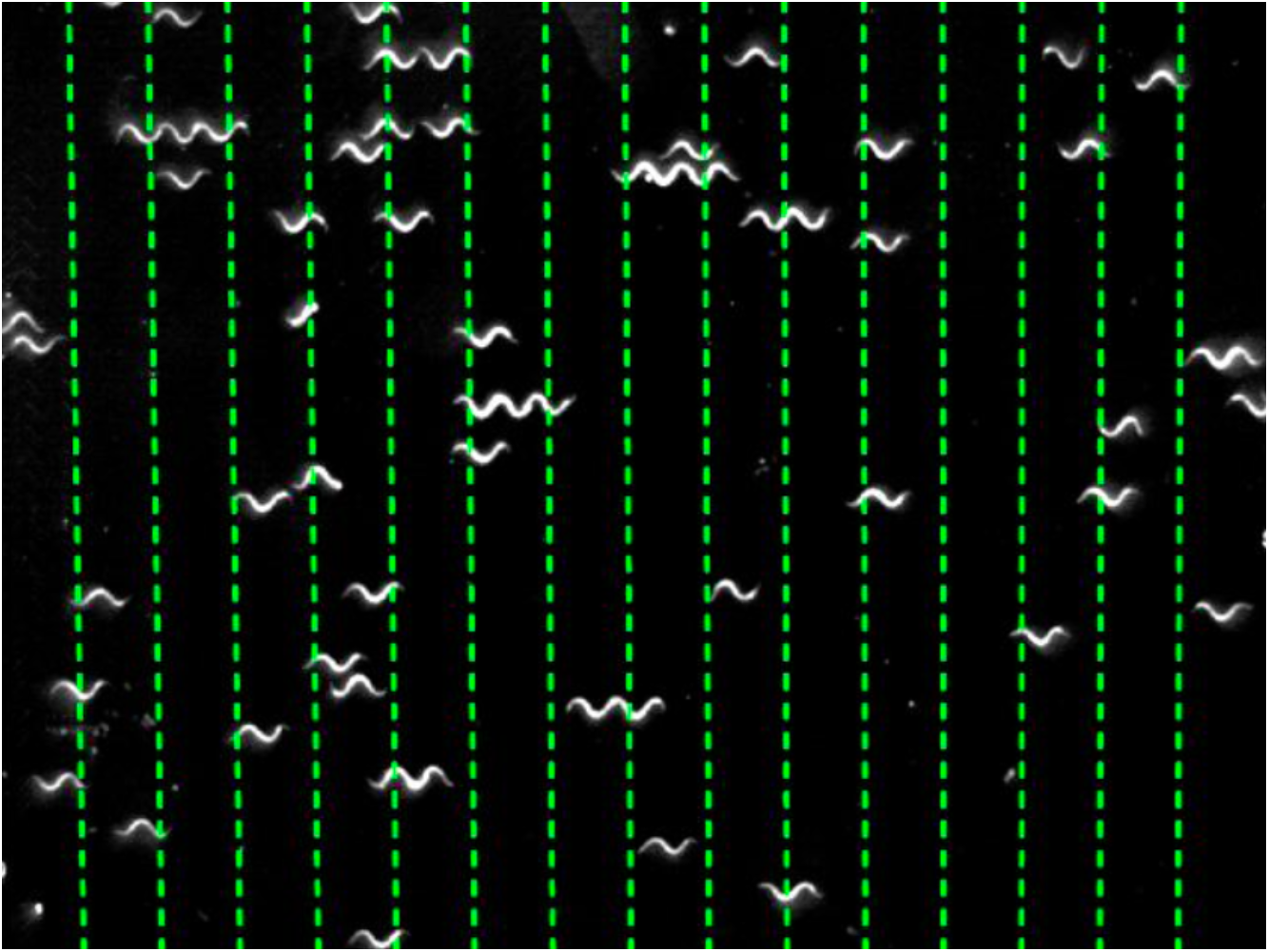
*C. elegans* responding to a gentle (30 μm) stimulus. Touch channel locations are shown in green, changing to red during stimulus pressurization. Field of view is 17.1 mm × 12.9 mm.

**Video S4:**
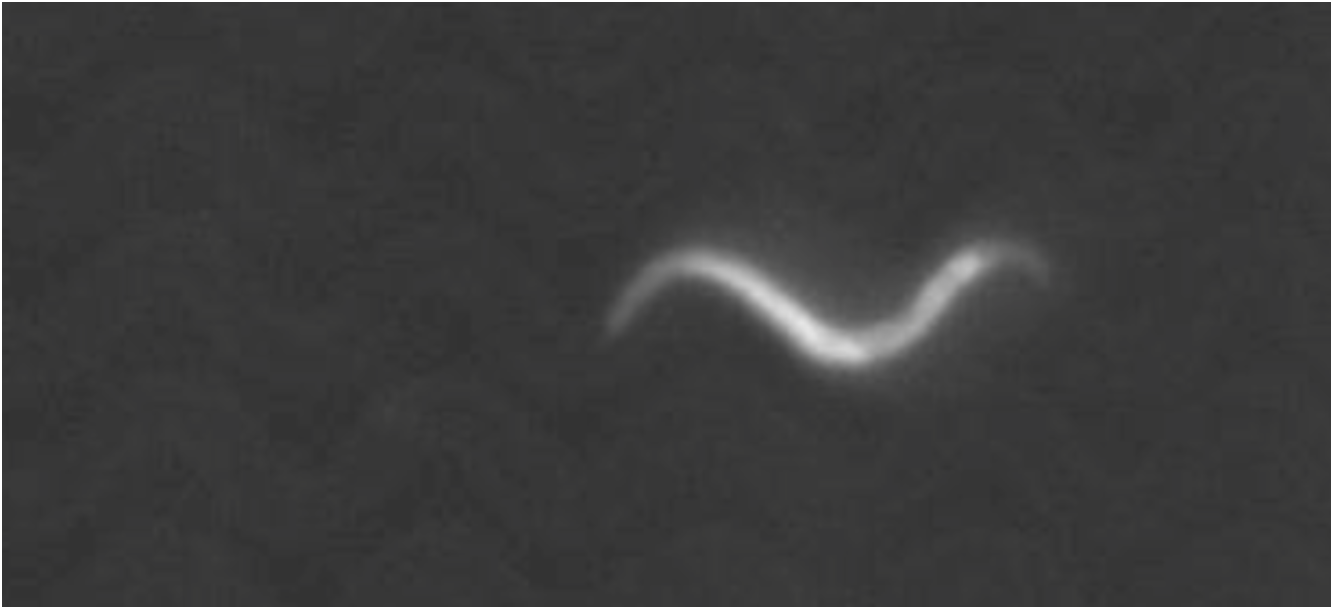
Detail of device showing *C. elegans* executing a 180 degree turn in a microfluidic channel. Field of view is 2.5 mm × 1.2 mm.

### S5 Details of force estimates

#### A. Estimate of forces during stimulation in microfluidic device

To estimate the force on a worm during touch stimulation, we first modeled the worm as a block of homogenous material with the elastic modulus of 125 kPa based on literature estimates for *C. elegans*^30,31^. We used a width and height of 75 μm based on the width of the worm, and a length of 100 μm to match the length of a touch valve. To compress this block 9.4 μm, the valve deflection resulting in a 50% response rate in WT worms, would require a force of 116 μN over an area of 7500 μm^2^.

A more realistic approach is to model the worm as a cylinder of diameter 75 μm and the touch valve ceiling as a plane applying pressure along a contact length of 100 μm and apply the following relationship^37^:

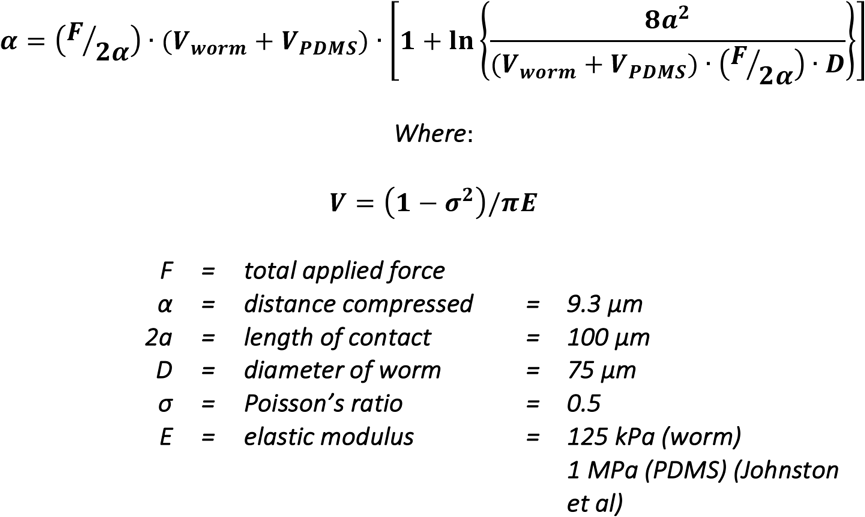

Assuming the above values and solving for the force gives a force of F = 94 μN, only slightly less than the simpler estimate.

#### B. Estimate of the pressure of the gentle touch response threshold

To estimate the pressure applied to the worm during a 50% threshold gentle (WT) touch stimulus, we divided these values by the estimated contact area projected in the direction of the applied force. For the stimulus applied in Petzold *et al*. a glass bead of diameter 10 μm pressed 1.5 μm into an approximately planar worm surface, this area would be a circle of 40 μm^2^, resulting in a pressure of 18.7 kPa. For our device, we assumed a force of 94 μN over a rectangular contact area of 49 μm × 100 μm, which would result from flattening of 75 μm wide cylindrical worm by 9.3 μm, resulting in a pressure of 19.0 kPa.

**Fig. S6.**
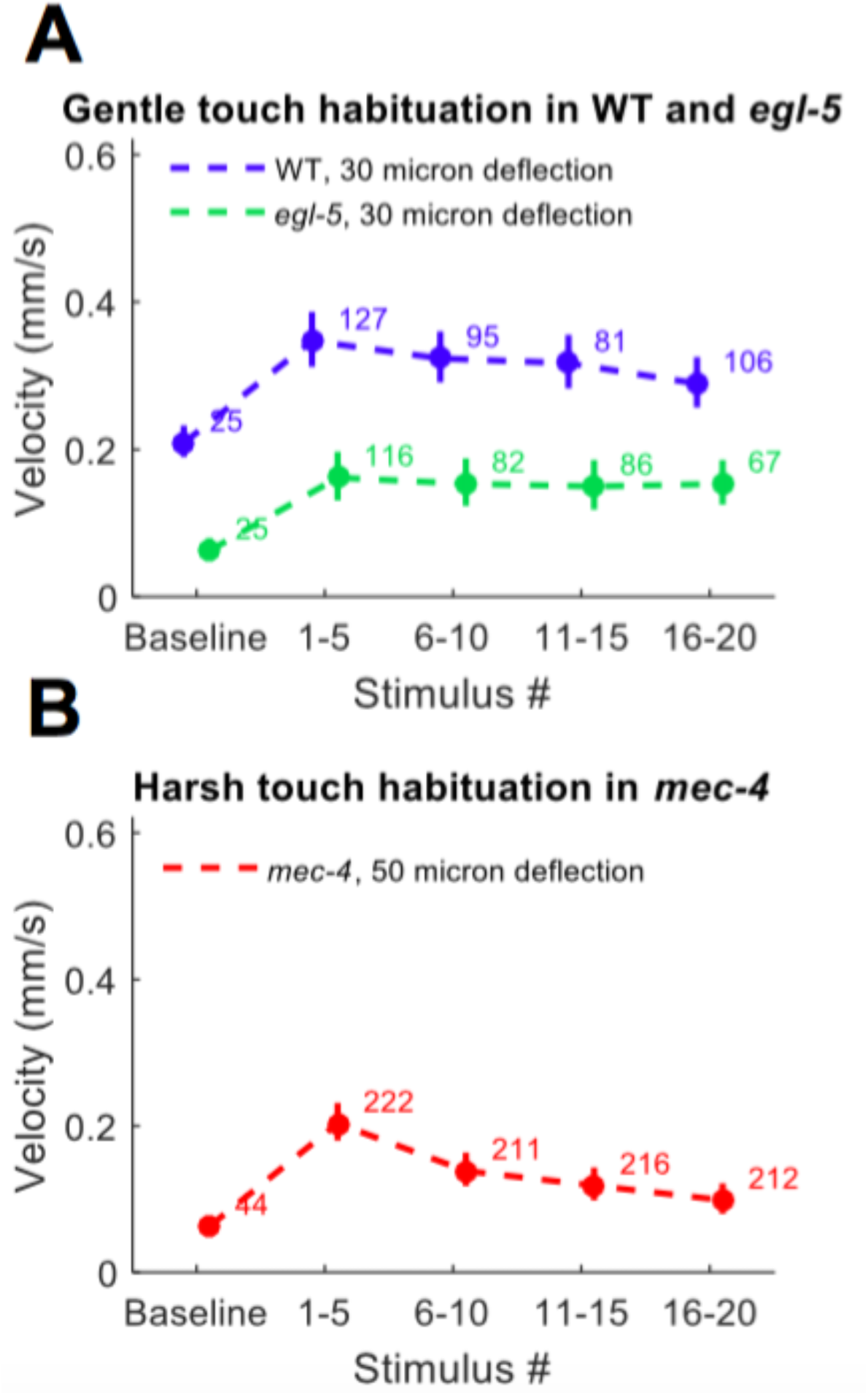
Velocity after repeated gentle (a) and harsh (b) touch stimulation. Average absolute value velocity ±SE of WT, *egl-5(u202)*, and *mec-4(u253)* animals to repeated gentle touch stimuli are binned into groups of five consecutive stimuli. The number of animals scored is shown for each point. Velocity changes are significantly higher than baseline, and the first five stimuli are not significantly different from the last five for gentle touch, but are for harsh touch. (Wilcoxon rank sum test, α = 0.05)

**Fig. S7.**
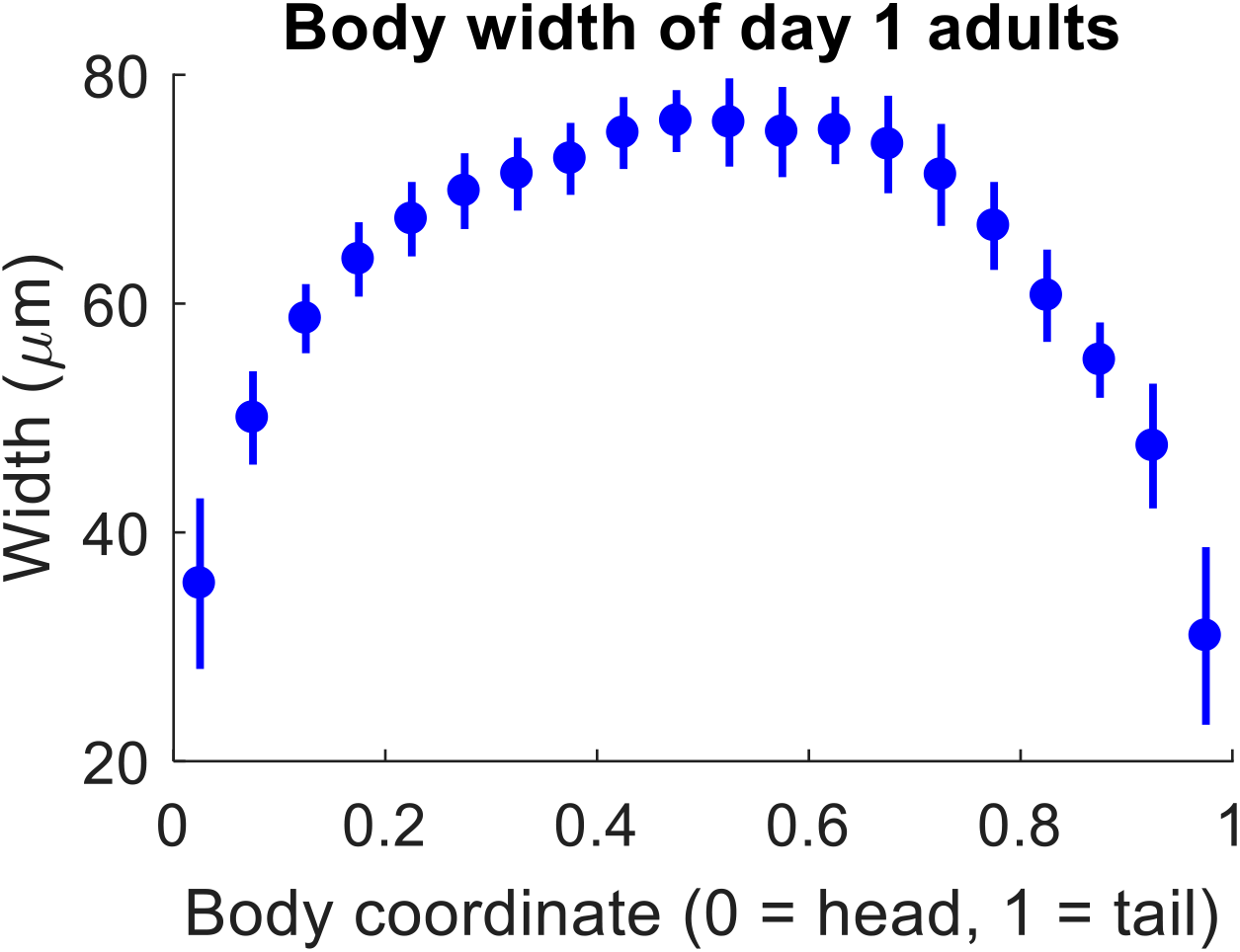
Width as a function of body coordinate for synchronized N2 day 1 adult animals, measured by analyzing micrographs of 10 individual worms. Measurements within each 5% of body length are binned together. Mean ± SD is shown.

**Fig. S8.**
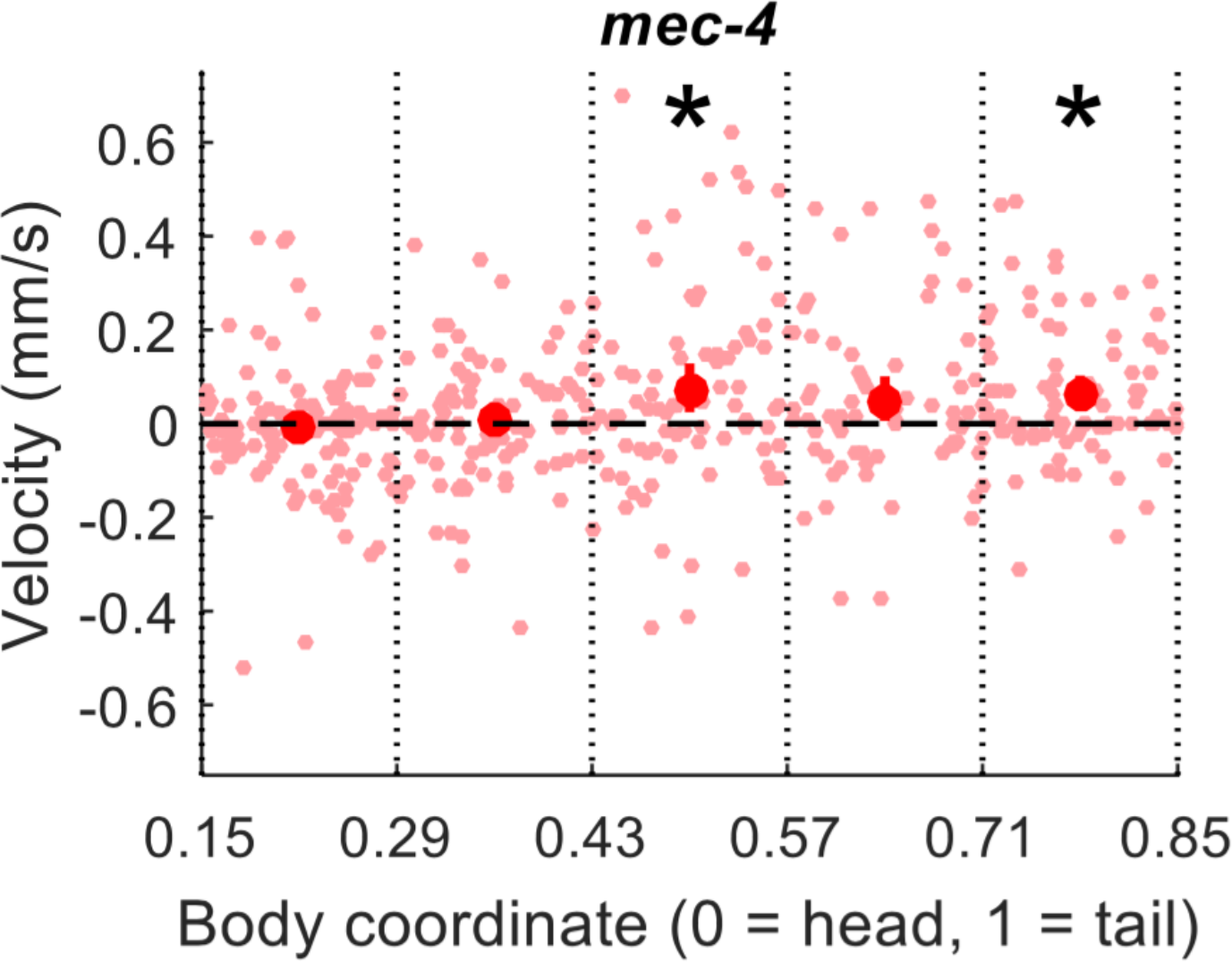
Gentle touch response field of *mec-4* mutants. Analysis and representation are as in Fig. 4.

## References

1. Chalfie, M. & Sulston, J. Developmental Genetics of the Mechanosensory Neurons of Caenorhabditis elegans. Dev. Biol. 82, 358–370 (1981).

2. Way, J.C., and Chalfie, M. The *mec-3* gene of Caenorhabditis elegans requires its own product for maintained expression and is expressed in three neuronal cell types. Genes Dev 3, 1823–1833 (1989).

3. Chatzigeorgiou, M. et al. Specific roles for DEG/ENaC and TRP channels in touch and thermosensation in C. elegans nociceptors. Nat. Neurosci. 13, 861–868 (2010).

4. Li, W., Kang, L., Piggott, B. J., Feng, Z. & Xu, X. Z. S. The neural circuits and sensory channels mediating harsh touch sensation in C. elegans. Nat. Commun. 1–22 (2011). doi:10.1038/ncomms1308

5. Chalfie, M. et al. The neural circuit for touch sensitivity in Caenorhabditis elegans. J. Neurosci. 5, 956–64 (1985).

6. Chalfie M., et al. Assaying mechanosensation (July 31, 2014), WormBook, ed. The *C. elegans* Research Community, WormBook, doi/10.1895/wormbook.1.172., http://www.wormbook.org.

7. Chen, X. & Chalfie, M. Modulation of C. elegans touch sensitivity is integrated at multiple levels. J. Neurosci. 34, 6522–36 (2014).

8. Chaudhuri, J. et al. A Caenorhabditis elegans Model Elucidates a Conserved Role for TRPA1-Nrf Signaling in Reactive α-Dicarbonyl Detoxification. Curr. Biol. 26, 3014–3025 (2016).

9. Suzuki, H. et al. In vivo imaging of C. elegans mechanosensory neurons demonstrates a specific role for the MEC-4 channel in the process of gentle touch sensation. Neuron 39, 1005–1017 (2003).

10. Chatzigeorgiou, M. et al. Specific roles for DEG/ENaC and TRP channels in touch and thermosensation in C. elegans nociceptors. Nat. Neurosci. 13, 861–8 (2010).

11. Julius, D. & Basbaum, A. I. Molecular mechanisms of nociception. Nature 413, 203–210 (2001).

12. Albeg, A. et al. C. elegans multi-dendritic sensory neurons: Morphology and function. Mol. Cell. Neurosci. 46, 308–317 (2011).

13. Brenner, S. The genetics of Caenorhabditis elegans. Genetics 77, 71–94 (1974).

14. Rankin, C. H., Beck, C. D. & Chiba, C. M. Caenorhabditis elegans: a new model system for the study of learning and memory. Behav. Brain Res. 37, 89–92 (1990).

15. Chiba, C. M. & Rankin, C. H. A developmental analysis of spontaneous and reflexive reversals in the nematode Caenorhabditis elegans. J. Neurobiol. 21, 543–554 (1990).

16. Ebrahimi, C. M. & Rankin, C. H. Early patterned stimulation leads to changes in adult behavior and gene expression in C. elegans. Genes, Brain Behav. 6, 517–528 (2007).

17. Wicks, S. R. & Rankin, C. H. Integration of mechanosensory stimuli in Caenorhabditis elegans. J. Neurosci. 15, 2434–2444 (1995).

18. Park, S.-J., Goodman, M. B. & Pruitt, B. L. Analysis of nematode mechanics by piezoresistive displacement clamp. Proc. Natl. Acad. Sci. U. S. A. 104, 17376–17381 (2007).

19. Petzold, B. C., Park, S.-J., Mazzochette, E. a, Goodman, M. B. & Pruitt, B. L. MEMS-based force-clamp analysis of the role of body stiffness in C. elegans touch sensation. Integr. Biol. (Camb). 5, 853–64 (2013).

20. Eastwood, A. L. et al. Tissue Mechanics Govern the Rapidly Adapting and Symmetrical Response to Touch. PNAS 113, E6955–E6963 (2016).

21. Nekimken, A. L. et al. Pneumatic stimulation of C. elegans mechanoreceptor neurons in a microfluidic trap. Lab Chip 17, 1116–1127 (2017).

22. Cho, Y. et al. Automated and controlled mechanical stimulation and functional imaging in vivo in C. elegans. Lab Chip (2017). doi:10.1039/C7LC00465F

23. O’Hagan, R., Chalfie, M. & Goodman, M. B. The MEC-4 DEG/ENaC channel of Caenorhabditis elegans touch receptor neurons transduces mechanical signals. Nat. Neurosci. 8, 43–50 (2005).

24. Hulme, S. E., Shevkoplyas, S. S., Apfeld, J., Fontana, W. & Whitesides, G. M. A microfabricated array of clamps for immobilizing and imaging C. elegans. Lab Chip 7, 1515 (2007).

25. Lockery, S. R. et al. Artificial dirt: microfluidic substrates for nematode neurobiology and behavior. J. Neurophysiol. 99, 3136–43 (2008).

26. Unger, M. A., Chou, H. P., Thorsen, T., Scherer, A. & Quake, S. R. Monolithic microfabricated valves and pumps by multilayer soft lithography. Science 288, 113–116 (2000).

27. Khanafer, K., Duprey, A., Schlicht, M. & Berguer, R. Effects of strain rate, mixing ratio, and stress-strain definition on the mechanical behavior of the polydimethylsiloxane (PDMS) material as related to its biological applications. Biomed. Microdevices 11, 503–508 (2009).

28. Wang, Z., Volinsky, A. A. & Gallant, N. D. Crosslinking effect on polydimethylsiloxane elastic modulus measured by custom-built compression instrument. J. Appl. Polym. Sci. 41050, 1–4 (2014).

29. Stiernagle, T. Maintenance of *C. elegans* (February 11, 2006), WormBook, ed. The *C. elegans* Research Community, WormBook, doi/10.1895/wormbook.1.101.1, http://www.wormbook.org.

30. Backholm, M., Ryu, W. S. & Dalnoki-Veress, K. Viscoelastic properties of the nematode Caenorhabditis elegans, a self-similar, shear-thinning worm. Proc. Natl. Acad. Sci. 110, 4528–4533 (2013).

31. Gilpin, W., Uppaluri, S. & Brangwynne, C. P. Worms under pressure: Bulk mechanical properties of C. elegans are independent of the cuticle. Biophys. J. 108, 1887–1898 (2015).

32. Johnston, I. D., McCluskey, D. K., Tan, C. K. L. & Tracey, M. C. Mechanical characterization of bulk Sylgard 184 for microfluidics and microengineering. J. Micromechanics Microengineering 24, 35017 (2014).

33. Sugi, T., Ohtani, Y., Kumiya, Y., Igarashi, R. & Shirakawa, M. High-throughput optical quantification of mechanosensory habituation reveals neurons encoding memory in Caenorhabditis elegans. PNAS 111, 17236–17241 (2014).

34. Takuma, S., Okumura, E., Kaori, K. & Igarashi, R. Nanoscale Mechanical Stimulation Method for Quantifying C. elegans Mechanosensory Behavior and Memory. Anal. Sci. 32, 1159–1164 (2016).

35. Timbers, T. a., Giles, A. C., Ardiel, E. L., Kerr, R. a. & Rankin, C. H. Intensity discrimination deficits cause habituation changes in middle-aged Caenorhabditis elegans. Neurobiol. Aging 34, 621–631 (2013).

36. Hart, A. C., ed. Behavior (July 3, 2006), WormBook, ed. The *C. elegans* Research Community, WormBook, doi/10.1895/wormbook.1.87.1, http://www.wormbook.org.

37. Puttock, M. J. & Thwaite, E. G. Elastic Compression of Spheres and Cylinders at Point and Line Contact. Natl. Stand. Lab. Tech. Pap. No. 25 1–64 (1969).

38. Chalfie, M. & Au, M. Genetic Control of Differentiation of the Caenorhabditis elegans Touch Receptor Neurons. Science (80-.). 243, 1027–1033 (1987).

39. Roberts, W. M. et al. A stochastic neuronal model predicts random search behaviors at multiple spatial scales in C. elegans. Elife 5, 1–30 (2016).

40. Suzuki, H. et al. In vivo imaging of C. elegans mechanosensory neurons demonstrates a specific role for the MEC-4 channel in the process of gentle touch sensation. Neuron 39, 1005–1017 (2003).

